# Yield is negatively correlated with nucleotide-binding leucine-rich repeat gene content in soybean

**DOI:** 10.1101/2021.12.12.472330

**Authors:** Philipp E. Bayer, Haifei Hu, Jakob Petereit, Mark C Derbyshire, Rajeev K. Varshney, Babu Valliyodan, Henry T. Nguyen, Jacqueline Batley, David Edwards

**Affiliations:** School of Biological Sciences, University of Western Australia, Perth, WA, Australia; Centre for Crop and Disease Management, School of Molecular and Life Sciences, Curtin University, Bentley, WA, 6102, Australia; State Agricultural Biotechnology Centre, Centre for Crop and Food Innovation, Food Futures Institute, Murdoch University, Murdoch, Western Australia, 6150 Australia; Center of Excellence in Genomics & Systems Biology, International Crops Research Institute for the Semi-Arid Tropics (ICRISAT), Hyderabad, Telangana, 502324 India; Division of Plant Sciences and National Center for Soybean Biotechnology, University of Missouri, Columbia, MO, 65211, USA; Department of Agriculture and Environmental Sciences, Lincoln University, Jefferson City, MO, 65101, USA

## Abstract

The availability of increasing quantities of crop pangenome data permits the detailed association of gene content with agronomic traits. Here, we investigate disease resistance gene content of diverse soybean cultivars and report a significant negative correlation between the number of NLR resistance (*R*) genes and yield. We find no association between *R-*genes with seed weight, oil or protein content, and we find no correlation between yield and the number of RLK, RLP genes, or the total number of genes. These results suggest that recent yield improvement in soybean may be partially associated with the selective loss of NLR genes. Three quarters of soybean NLR genes do not show presence/absence variation, limiting the ability to select for their absence, and so the deletion or disabling of select NLR genes may support future yield improvement.

## Main

Soybean is the leading legume crop globally, covering an estimated 6% of arable land ^1^. With advances in breeding and agronomy, soybean yield has increased from 15 kg ha^-1^ yr^-1^ prior to the 1980s, to 30 kg ha^-1^ yr^-1^ in the late 90s ^2^. However, since 2000, increases in soybean yield have been limited ^3^, and new strategies are needed to increase yield to meet the global demand for this crop. The application of genomics has the potential to accelerate the production of high yielding varieties adapted to a changing climate ^4^. Recent advances in pangenomics supports the study of gene presence/absence variation during selection ^5,6^. Pangenomes have been constructed in several crop species, including *Brassica oleracea* ^7,8^, soybean ^9-11^, sesame ^12^, wheat ^13^, and banana ^14^. Several genes demonstrating presence/absence variation influence crop traits, for example flavour in tomato ^15,16^, or submergence tolerance in rice ^17-19^. Dispensable genes are enriched for functions related to disease resistance ^8,10,20^ allowing breeders to select for their presence or absence.

We have previously identified an association between gene content and breeding in soybean ^10^. There is evidence for a continuing trade-off between disease resistance and yield ^21^. For example, evidence in *Arabidopsis* suggests a negative impact of additional disease resistance genes on yield ^22^, while in wheat there is a significant positive association between yield and disease resistance ^23^. We therefore examined the association between the abundance of candidate disease resistance gene classes and agronomic traits in soybean, including yield, seed weight, oil and protein content. We identified 486 nucleotide-binding domain leucine-rich repeat (NLR), 1,173 receptor-like kinase (RLK), and 180 receptor-like protein (RLP) gene candidates in the soybean pangenome (Supplementary Table 1), and of these three classes, 122 (25%), 89 (8%), and 45 (25%) NLRs, RLKs, and RLPs respectively were lost in at least one individual.

**Table 1:** Calculated slope for the number of NLR genes and the intercept for the mixed-effect model using landrace as the baseline (**: p<0.005 using t-tests with Satterthwaite’s method)

| Predictors | Estimates | 95% CI | p-value |
| --- | --- | --- | --- |
| Intercept (Country) | 9.307 | 4.295 – 14.318 | **<0.001 \*\*** |
| NLR count | -0.016 | -0.027 - -0.005 | **0.005 \*\*** |
| Group Old cultivar | -0.218 | -0.491 – 0.056 | 0.118 |
| Group Modern cultivar | 0.801 | 0.380 – 1.222 | **<0.001 \*\*** |

We investigated whether loss of disease resistance genes is associated with agronomic traits, especially yield, in soybean (Supplementary Table 2). We hypothesised that some of the previously observed overall gene loss may have been driven by breeders selecting against costly NLR genes. As expected, NLRs showed a slight reduction in number during domestication from wild *G. soja* to landraces (average 8 genes lost, Mann-Whitney U test, p < 0.001) and during breeding from old to modern cultivars (average 3 genes lost, Mann-Whitney U test, p < 0.001, Figure 1). Similar results have been observed in rice, where the genome of the wild ancestor *Oryza rufipogon* contained between 2% and 14% more NLR genes than three of four domesticated rice genome assemblies ^24^.

**Figure 1:** Reduction of NLR gene content across the history of soybean breeding (***: p < 0.001, **: p < 0.01, Mann-Whitney U test) showing a reduction of NLR content in two steps: once during domestication, and once during the breeding of modern cultivars.

**Figure 1:** Linear model of yield based on the number of NLR genes, accession group, and country of origin) showing a statistically significant association between yield and the number of NLR genes). The plot is coloured by country with the five most-common countries labelled.

We found a significant difference in RLKs and RLPs during domestication between *G. soja* and landraces. We found no such association during modern breeding with one gene lost on average from wild *G. soja* to landraces, and no genes lost on average from old to modern cultivars (Mann-Whitney U test p > 0.05 in both cases) (Supplementary Figures 1, 2).

To further investigate whether the loss of NLR genes in modern cultivars is driven by yield and not by other breeding targets, we examined associations between yield and NLR numbers using linear regression while accounting for accession groups and country of origin. We assumed that different accession groups (landrace, old cultivar, modern cultivar) and the accession’s country of origin had an impact on the trajectory of gene loss due to different aims of different breeders, so we included accession groups and country of origin as covariates in the model linking NLR count with yield. Linear modelling revealed a significant negative correlation between the number of NLR genes and yield (−0.02, p < 0.005, t = -3.5, Figure 2, Table 3, Supplementary Figures 3-5).

**Figure 2:** Predicted values of non-normalised yield using the NLR count based on the linear model.

Visualising the model’s predicted yield by NLR count shows from 465 to 430 NLR genes, predicted yield rises by an average of 0.58 Mg ha^-1^ (Figure 3). In modern cultivars, assuming a yield range of 3.1 to 2.5 Mg ha^-1^, each additional NLR gene has an average yield penalty of 1.6%.

**Figure 3:**
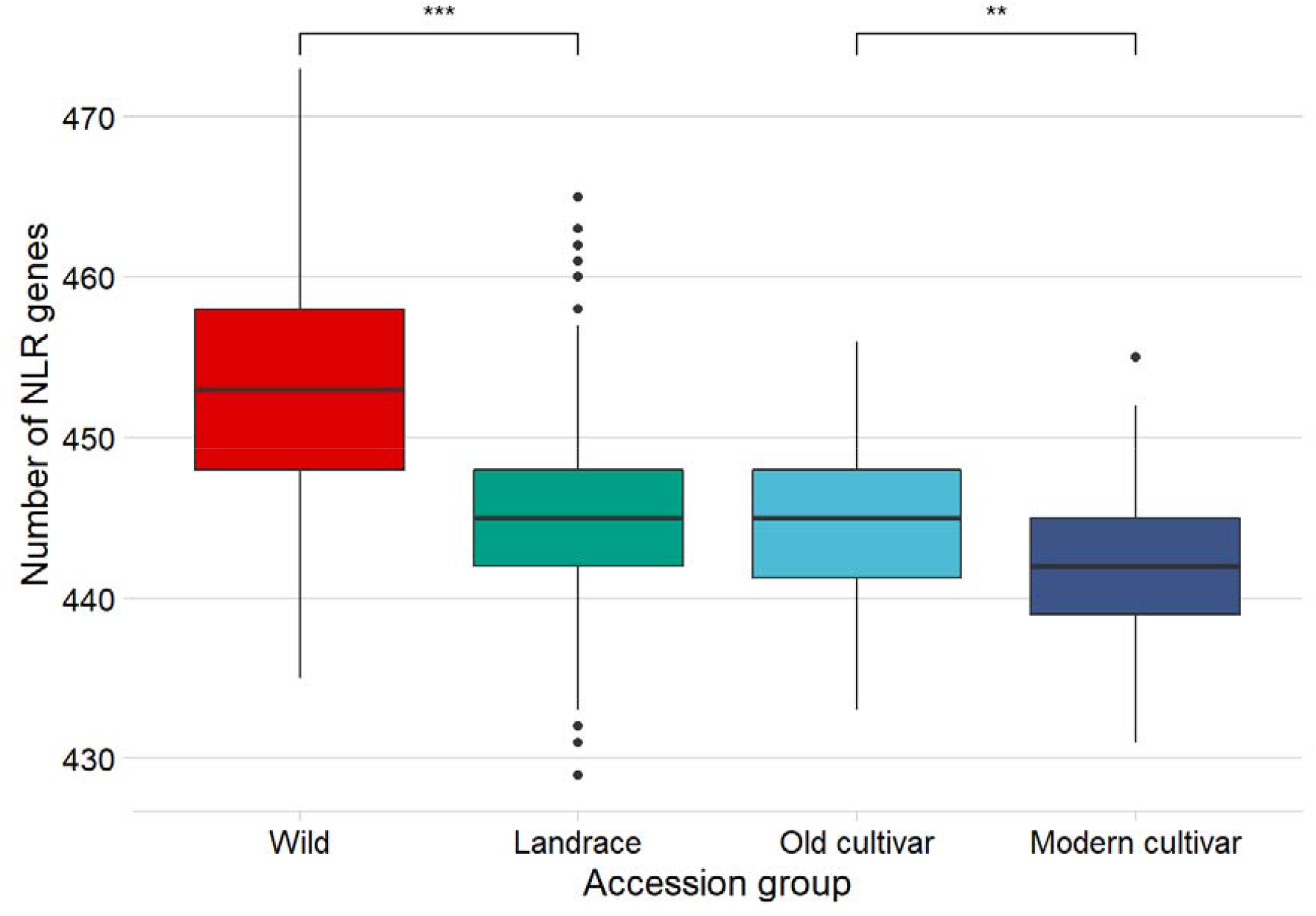
Reduction of NLR gene content across the history of soybean breeding (***: p < 0.001, **: p < 0.01, Mann-Whitney U test) showing a reduction of NLR content in two steps: once during domestication, and once during the breeding of modern cultivars.

**Figure 4:**
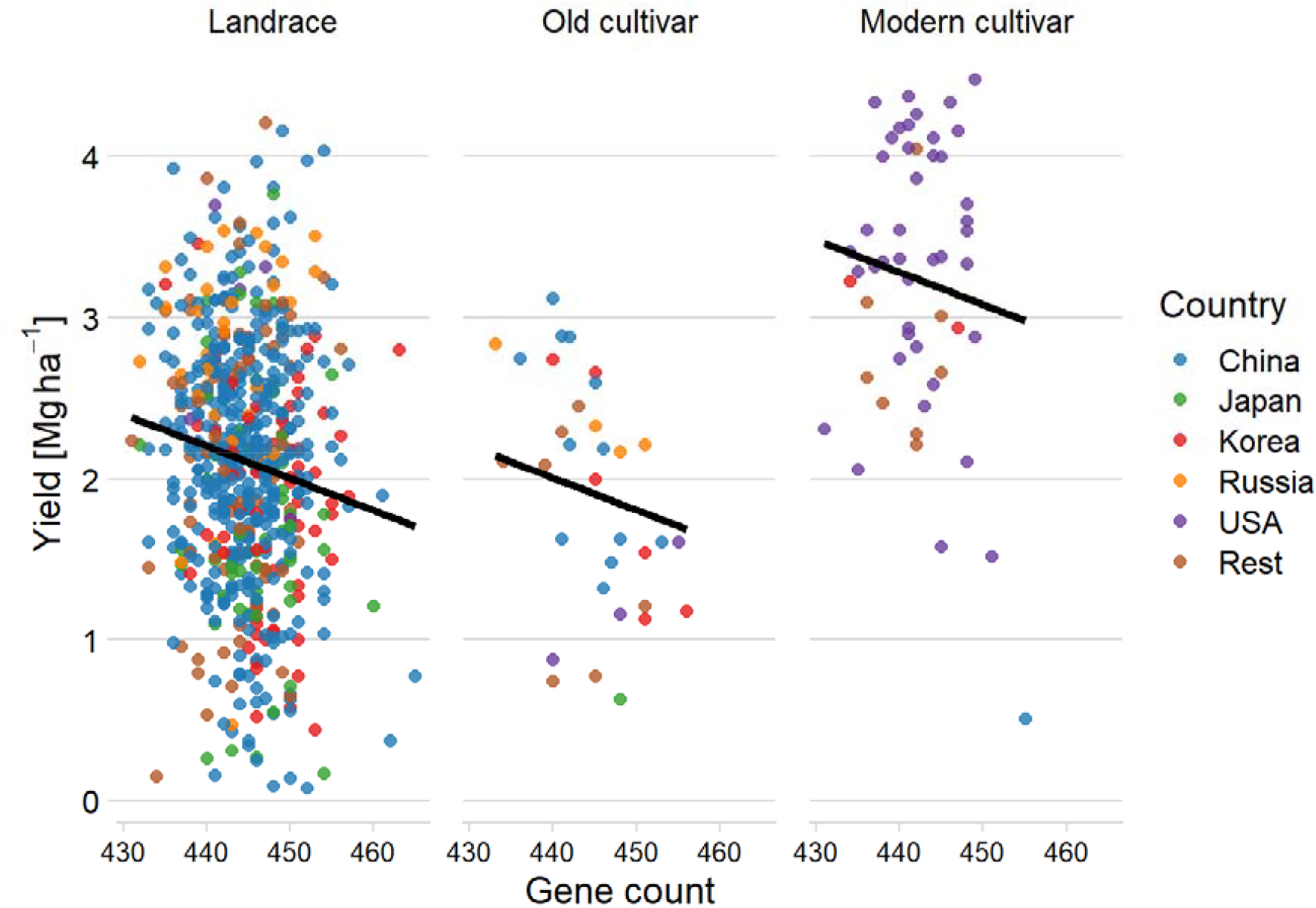
Linear model of yield based on the number of NLR genes, accession group, and country of origin showing a statistically significant association between yield and the number of NLR genes). The plot is coloured by country with the five most-common countries labelled. Predictions of the model for all countries averaged are shown in black.

**Figure 5:**
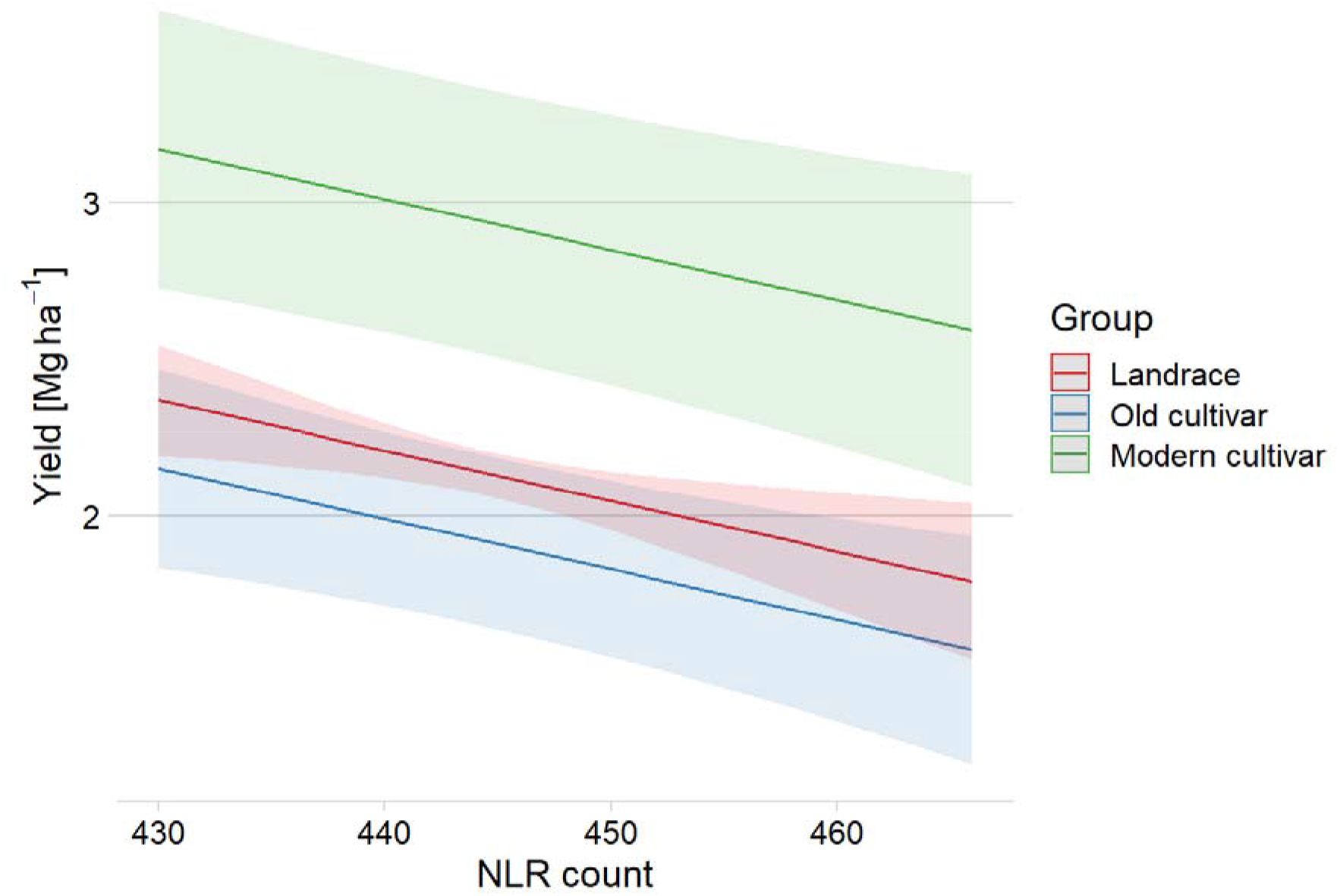
Predicted interaction effect on yield by the number of NLR genes by group

We searched for NLR genes directly linked with yield to learn whether the difference in yield could be attributed to single NLR genes with strong impacts. To this end we carried out a genome-wide association study linking presence/absence in NLR genes with yield (PAV-GWAS). We found four candidate gene associations with FDR-adjusted p-value < 0.05. One NL gene was located on the reference genome assembly (GlymaLee.01G030900), and three NBS genes were assembled during pangenome construction (UWASoyPan00316, UWASoyPan00772, UWASoyPan04354, Supplementary Table 3). These genes show different impacts on yield with landraces carrying the genes GlymaLee.01G030900 and UWASoyPan00772 showing higher yield, while landraces carrying the genes UWASoyPan00316 showed lower yield, with no statistically significant differences in individuals carrying UWASoyPan04354 (Supplementary Figure 6). Landraces carrying GlymaLee.01G030900 demonstrated an average increased yield of 0.22 Mg ha^-1^, contrary to the trend of negative association of NLR genes. In contrast, landraces carrying UWASoyPan00316 showed an average lower yield of 0.20 Mg ha^-1^ (Supplementary Table 4). The presence of the four candidate genes in the population shows a generally negative trend during domestication and breeding from an average of 67.9% in *G. soja* to an average of 33.7% presence in modern cultivars. The only PAV-GWAS gene that appears more often in modern cultivars than old cultivars, contrary to the overall pattern of fewer NLR-genes, is GlymaLee.01G030900, which is present in 71% of modern cultivars and just 35% of old cultivars. However, the yield changes associated with the four candidate genes do not explain all the difference in yield observed.

To examine whether the link between NLR genes and yield was not due to the previously observed overall gene loss in soybean during domestication and breeding ^10^, we examined whether there was an association between overall gene content and yield. We found no significant association between the number of genes per line and yield when accounting for accession group and country of origin (−0.00004, p = 0.8, t = -0.2, Supplementary Figure 8, Supplementary Table 5). This confirms that the negative association between NLR genes and yield is not due to the overall gene loss during soybean breeding. We found no significant association between NLR gene count and protein content, oil content, or seed weight (Supplementary Table 6). For these models, the association between country and the respective phenotype was always significant reflecting selection for these traits during breeding. We also found no significant association for RLPs and RLKs with yield. RLPs and RLKs typically have many other functions, for example, fourteen *Arabidopsis* genes are involved in cold-response regulation ^25^, so loss of these genes can have negative results unrelated to a loss of a specific disease resistance.

The association between NLR gene content and yield may be due to several factors. Activation of immunity pathways redirects hormone signalling away from plant growth ^21^ Another explanation is the potential cost of resistance gene expression, priming the plant for defence and reducing vigour and subsequent yield. In *Arabidopsis*, disease resistance gene mutation can lead to autoimmunity and cell death ^26-29^. Plants with a point mutation in *snc1*, a TNL NLR gene, over-accumulate the SNC1 protein leading to negative autoimmune responses ^30,31^. At least one pair of NLR-variants interacts in *Arabidopsis* triggering autoimmunity through the same pathway of plant NLRs recognising foreign elicitors ^26^. The introduction of NLR genes can also lead to a reduction in yield. For example, in *Arabidopsis*, transgenic introduction of the NLR *RPM1* led to a 9% decrease in total seed number in the absence of the pathogen ^22^.

Our results may explain why disease resistance genes are the most common class of genes among variable genes in many of the crop pangenomes assembled to date ^8,10,20,32^. In the *B. oleracea* pangenomes, 42% of predicted resistance genes were lost in at least one line compared to 19% of all *B. oleracea* genes ^7,8^, while in the amphidiploid *B. napus*, 69% of predicted resistance genes were lost in at least one line compared to 38% of all *B. napus* genes ^20,33^. If the presence of NLR-genes impacts yield across different crops, the identification and removal of NLR genes may provide a new avenue to improve crop yield using genomic tools such as CRISPR-Cas9 ^34^.

## Methods

The pangenome annotation of ^10^ available at https://doi.org/10.26182/5f34ac3377313 was searched for *R*-gene-specific protein domains and assigned to resistance gene candidate classes using RGAugury ^35^ v2016-11-10. Per-line *R-*gene counts were calculated using R ^36^ v3.6.3 and tidyverse ^37^ v1.3.0.

Phenotype data for all soybean lines was downloaded from GRIN-Global, U.S. National Plant Germplasm System. Yield was measured for 656 soybean landraces, 33 old and 52 modern cultivars. Oil and protein content were measured for 677 landraces, 41 old and 71 modern cultivars. Seed weight was measured for 548 landraces, 31 old and 54 modern cultivars.

The linear model was fit using lm() of R v3.6.3 using the formula *Yield ∼ NLR-count + Group + Country*. The resulting model was investigated using sjPlot ^38^ v2.8.6, lmerTest ^39^ v3.1-2, and ggeffects ^40^ v0.16.0. Models for other phenotypes were implemented using the same formula of *∼ NLR-count + Group + Country*.

The GWAS was carried out using GAPIT3 v3.1.0 ^41^ using Mixed Linear Model (MLM), Multiple Loci Mixed Model (MLMM), General Linear Model (GLM), and FarmCPU ^42^. We calculated principal components (PCs) based on the publicly available SNP data from ^10^ using SNPRelate v 1.20.1 ^43^ with LD-pruning set to 0.2, and used PC1 and PC2 as covariates in GAPIT instead of GAPIT’s in-built function. Genes lost in fewer than 5% of individuals were removed from the analysis.

The analysis is fully reproducible using R ^36^ v3.6.3, tidyverse ^37^ v1.3.0, and workflow ^44^ v1.6.2.9000 using the code and data available at https://philippbayer.github.io/R_gene_analysis/ and https://github.com/philippbayer/R_gene_analysis.

## Acknowledgments

This work is funded by the Australian Research Council (Projects DP1601004497, LP160100030, DP210100296, DP200100762, and DE210100398). This work was supported by resources provided by the Pawsey Supercomputing Centre with funding from the Australian Government and the Government of Western Australia. Soybean data was generated by the funding support to H.N and B.V. from the United Soybean Board (U.S.A) and former Bayer Crop Science, Dow AgroSciences, and BASF (project #1320-532-5615). Funding from the USDA-NIFA Allen to B.V. (project #1020002) is acknowledged. P. E. B. thanks Dr. Giovanni Polverino and Dr. Lyron Winderbaum for their valuable feedback on the modelling results.

## Author contributions

P.E.B., H.H., and M.C.D carried out the analysis. B. V., H. N., and R.J.K. supplied data. P.E.B., M.C.D., B. V., H.N, J.B. and D.E. wrote the manuscript.

## References

1 Valliyodan, B. et al. Genetic diversity and genomic strategies for improving drought and waterlogging tolerance in soybeans. Journal of experimental botany 68, 1835–1849 (2017).

2 Specht, J., Hume, D. & Kumudini, S. Soybean yield potential—a genetic and physiological perspective. Crop science 39, 1560–1570 (1999).

3 Ray, D. K., Ramankutty, N., Mueller, N. D., West, P. C. & Foley, J. A. Recent patterns of crop yield growth and stagnation. Nature communications 3, 1–7 (2012).

4 Abberton, M. et al. Global agricultural intensification during climate change: a role for genomics. Plant biotechnology journal 14, 1095–1098 (2016).

5 Golicz, A. A., Batley, J. & Edwards, D. Towards plant pangenomics. Plant Biotechnology Journal 14, 1099–1105, doi:10.1111/pbi.12499 (2016).

6 Bayer, P., Golicz, A., Scheben, S., Batley, J. & Edwards, D. Plant pan-genomes are the new reference. Nature Plants (2020).

7 Golicz, A. A. et al. The pangenome of an agronomically important crop plant Brassica oleracea. Nature Communications 7, 13390, doi:10.1038/ncomms13390 (2016).

8 Bayer, P. E. et al. Variation in abundance of predicted resistance genes in the Brassica oleracea pangenome. Plant Biotechnology Journal 17, 789–800, doi:10.1111/pbi.13015 (2019).

9 Liu, Y. et al. Pan-Genome of Wild and Cultivated Soybeans. Cell 182, 162-176.e113, doi:10.1016/j.cell.2020.05.023 (2020).

10 Bayer, P. E. et al. Sequencing the USDA core soybean collection reveals gene loss during domestication and breeding. The Plant Genome, Art. e20109 (2021).

11 Torkamaneh, D., Lemay, M. A. & Belzile, F. The pan-genome of the cultivated soybean (PanSoy) reveals an extraordinarily conserved gene content. Plant Biotechnology Journal (2021).

12 Yu, J. et al. Insight into the evolution and functional characteristics of the pan-genome assembly from sesame landraces and modern cultivars. Plant Biotechnology Journal 17, 881–892 (2019).

13 Montenegro, J. D. et al. The pangenome of hexaploid bread wheat. The Plant Journal 90, 1007–1013 (2017).

14 Rijzaani, H. et al. The pangenome of banana highlights differences between genera and genomes. The Plant Genome n/a, e20100, doi:https://doi.org/10.1002/tpg2.20100 (2021).

15 Gao, L. et al. The tomato pan-genome uncovers new genes and a rare allele regulating fruit flavor. Nature Genetics 51, 1044–1051, doi:10.1038/s41588-019-0410-2 (2019).

16 Alonge, M. et al. Major Impacts of widespread structural variation on gene expression and crop improvement in tomato. Cell 182, 145-161. e123 (2020).

17 Xu, K. et al. Sub1A is an ethylene-response-factor-like gene that confers submergence tolerance to rice. Nature 442, 705–708 (2006).

18 Hattori, Y. et al. The ethylene response factors SNORKEL1 and SNORKEL2 allow rice to adapt to deep water. Nature 460, 1026–1030, doi:10.1038/nature08258 (2009).

19 Zhao, Q. et al. Pan-genome analysis highlights the extent of genomic variation in cultivated and wild rice. Nature Genetics 50, 278–284, doi:10.1038/s41588-018-0041-z (2018).

20 Dolatabadian, A. et al. Characterization of disease resistance genes in the Brassica napus pangenome reveals significant structural variation. Plant biotechnology journal (2019).

21 Ning, Y., Liu, W. & Wang, G.-L. Balancing immunity and yield in crop plants. Trends in plant science 22, 1069–1079 (2017).

22 Tian, D., Traw, M., Chen, J., Kreitman, M. & Bergelson, J. Fitness costs of R-gene-mediated resistance in Arabidopsis thaliana. Nature 423, 74–77 (2003).

23 Voss-Fels, K. P. et al. Breeding improves wheat productivity under contrasting agrochemical input levels. Nature plants 5, 706–714 (2019).

24 Stein, J. C. et al. Genomes of 13 domesticated and wild rice relatives highlight genetic conservation, turnover and innovation across the genus Oryza. Nature genetics 50, 285–296 (2018).

25 Lee, B.-h., Henderson, D. A. & Zhu, J.-K. The Arabidopsis cold-responsive transcriptome and its regulation by ICE1. The Plant Cell 17, 3155–3175 (2005).

26 Tran, D. T. et al. Activation of a plant NLR complex through heteromeric association with an autoimmune risk variant of another NLR. Current Biology 27, 1148–1160 (2017).

27 Rodriguez, E. et al. DNA damage as a consequence of NLR activation. PLoS genetics 14, e1007235 (2018).

28 Wan, W. L., Kim, S. T., Castel, B., Charoennit, N. & Chae, E. Genetics of autoimmunity in plants: an evolutionary genetics perspective. New Phytologist 229, 1215–1233 (2021).

29 Li, L. & Weigel, D. One Hundred Years of Hybrid Necrosis: Hybrid Autoimmunity as a Window into the Mechanisms and Evolution of Plant–Pathogen Interactions. Annual Review of Phytopathology 59 (2021).

30 Li, X., Clarke, J. D., Zhang, Y. & Dong, X. Activation of an EDS1-mediated R-gene pathway in the snc1 mutant leads to constitutive, NPR1-independent pathogen resistance. Molecular plant-microbe interactions 14, 1131–1139 (2001).

31 Zhang, Y., Goritschnig, S., Dong, X. & Li, X. A gain-of-function mutation in a plant disease resistance gene leads to constitutive activation of downstream signal transduction pathways in suppressor of npr1-1, constitutive 1. The Plant Cell 15, 2636–2646 (2003).

32 Inturrisi, F. et al. Genome-wide identification and comparative analysis of resistance genes in Brassica juncea. Molecular Breeding 40, 1–14 (2020).

33 Hurgobin, B. et al. Homoeologous exchange is a major cause of gene presence/absence variation in the amphidiploid Brassica napus. Plant Biotechnology Journal 16, 1265–1274 (2018).

34 Scheben, A., Wolter, F., Batley, J., Puchta, H. & Edwards, D. Towards CRISPR/Cas crops– bringing together genomics and genome editing. New Phytologist 216, 682–698 (2017).

35 Li, P. et al. RGAugury: a pipeline for genome-wide prediction of resistance gene analogs (RGAs) in plants. BMC genomics 17, 852 (2016).

36 R Core Team. R: A language and environment for statistical computing. (2020).

37 Wickham, H. et al. Welcome to the Tidyverse. Journal of Open Source Software 4, 1686 (2019).

38 Lüdecke, D. sjPlot: Data visualization for statistics in social science. R package version 2 (2018).

39 Kuznetsova, A., Brockhoff, P. B. & Christensen, R. H. lmerTest package: tests in linear mixed effects models. Journal of statistical software 82, 1–26 (2017).

40 Lüdecke, D. ggeffects: Tidy data frames of marginal effects from regression models. Journal of Open Source Software 3, 772 (2018).

41 Wang, J. & Zhang, Z. GAPIT Version 3: Boosting Power and Accuracy for Genomic Association and Prediction. bioRxiv (2020).

42 Liu, X., Huang, M., Fan, B., Buckler, E. S. & Zhang, Z. Iterative usage of fixed and random effect models for powerful and efficient genome-wide association studies. PLoS genetics 12, e1005767 (2016).

43 Zheng, X. et al. A high-performance computing toolset for relatedness and principal component analysis of SNP data. Bioinformatics 28, 3326–3328 (2012).

44 Blischak, J. D., Carbonetto, P. & Stephens, M. Creating and sharing reproducible research code the workflowr way. F1000Research 8 (2019).

