## Supplementary Figures for "Yield is negatively correlated with nucleotide-binding leucine-rich repeat gene content in soybean"

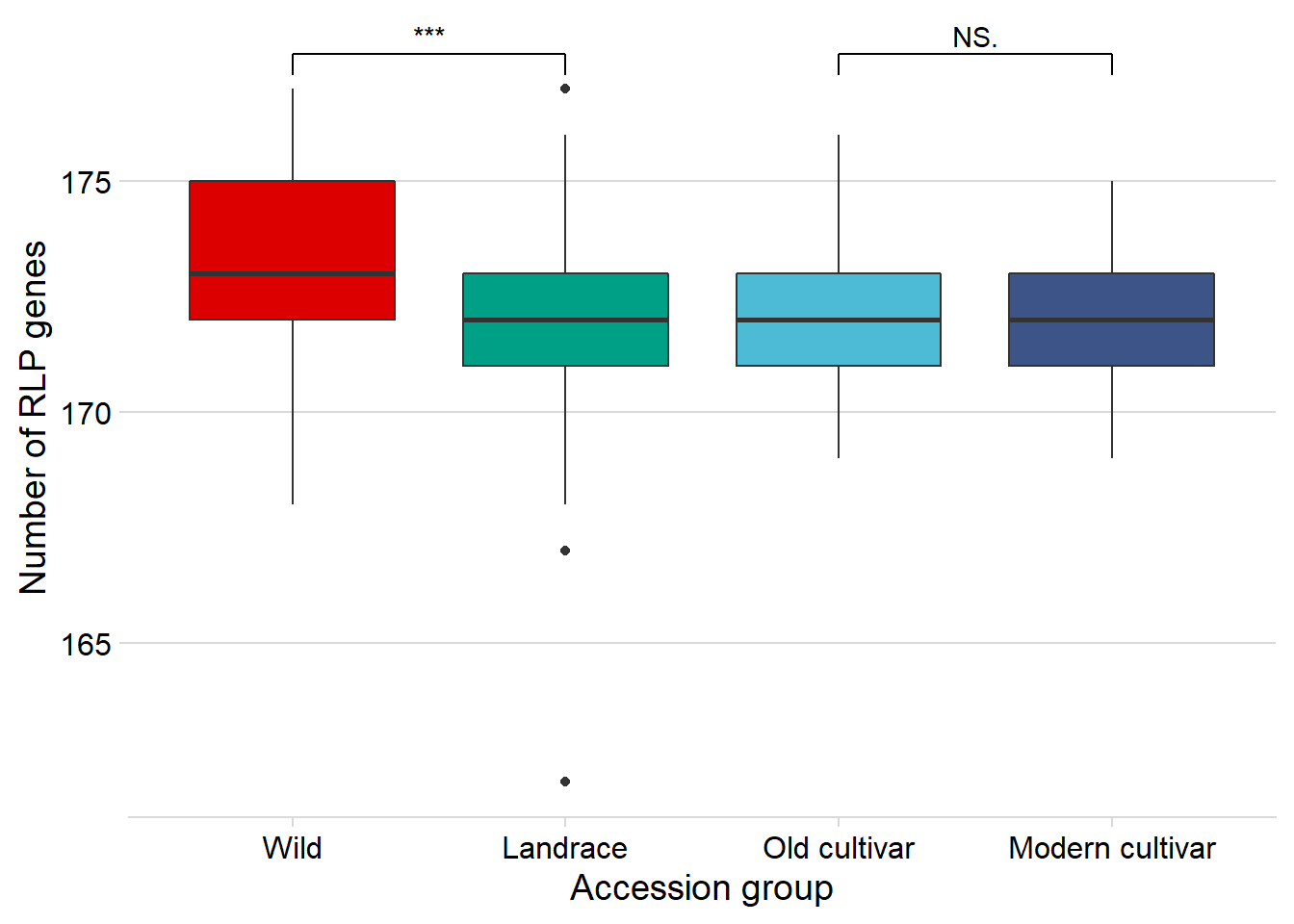

Supplementary Figure 1: Reduction of RLP gene content across the history of soybean breeding (***: p < 0.001, NS: not significant, Mann-Whitney U test) showing a reduction of RLP gene content only during domestication.

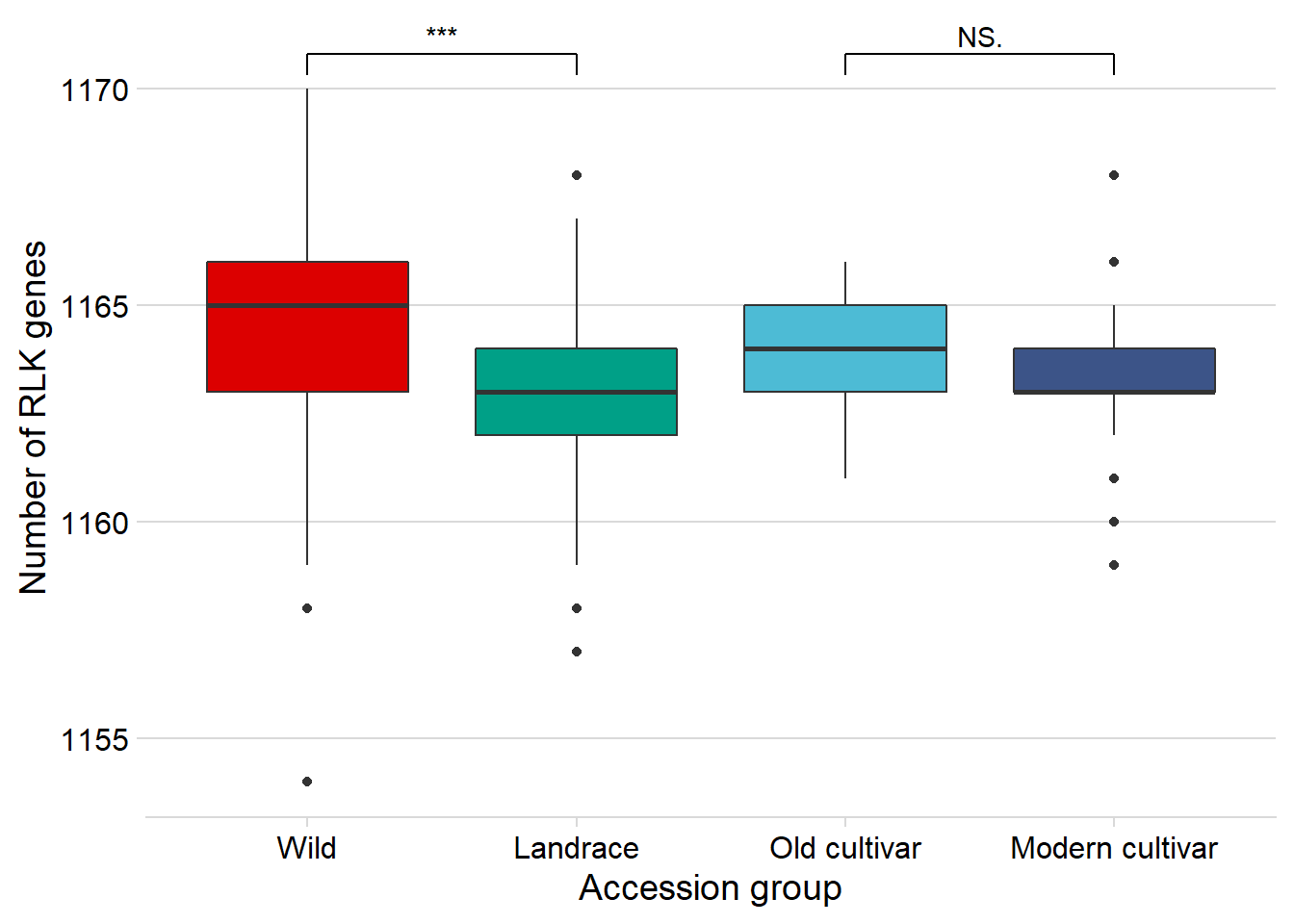

Supplementary Figure 2: Reduction of RLK gene content across the history of soybean breeding (***: p < 0.001, NS: not significant, Mann-Whitney U test) showing a reduction of RLK gene content only during domestication.

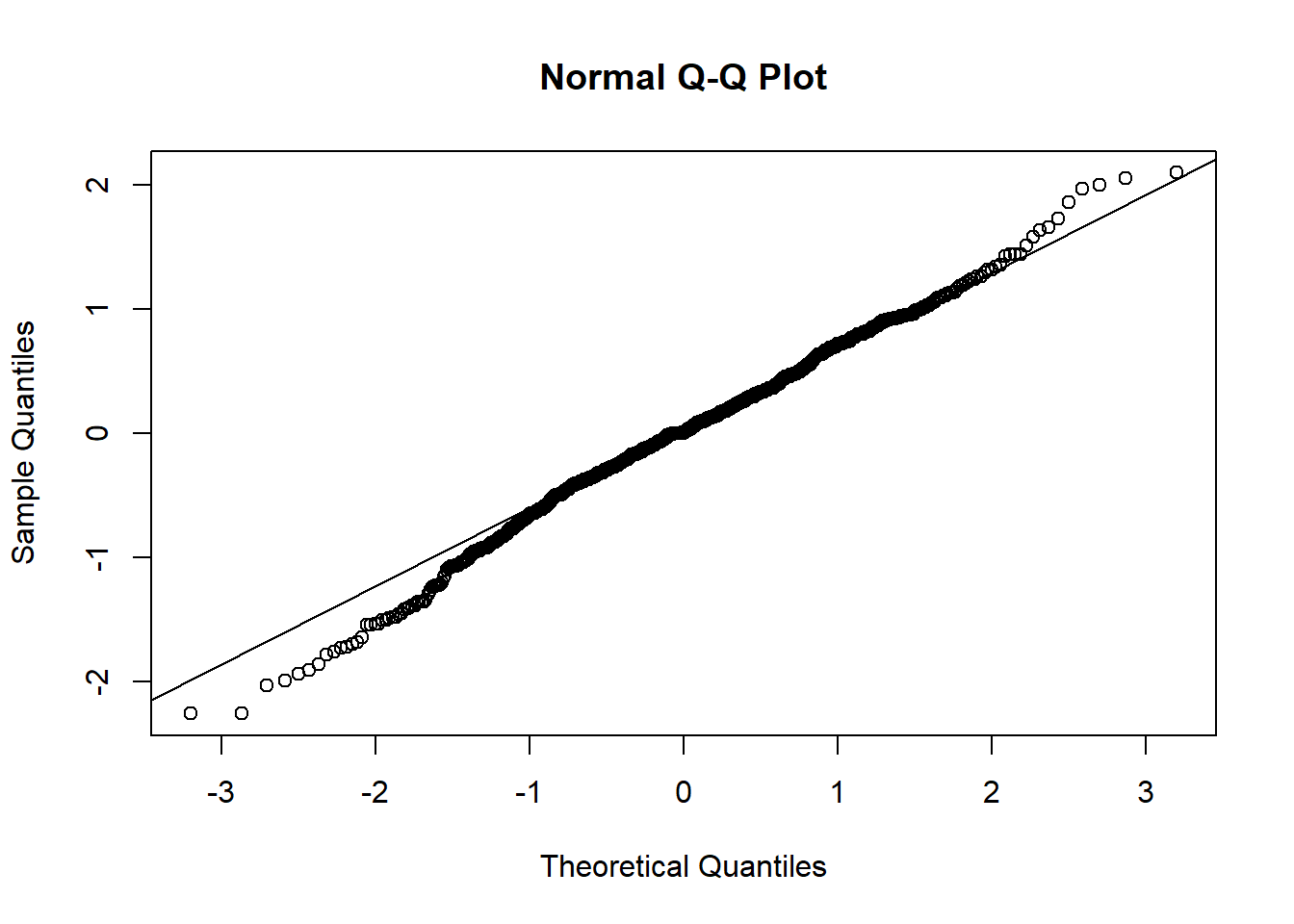

Supplementary Figure 3: Quantile-quantile plot for the linear model showing reasonable overlap between samples and the model prediction.

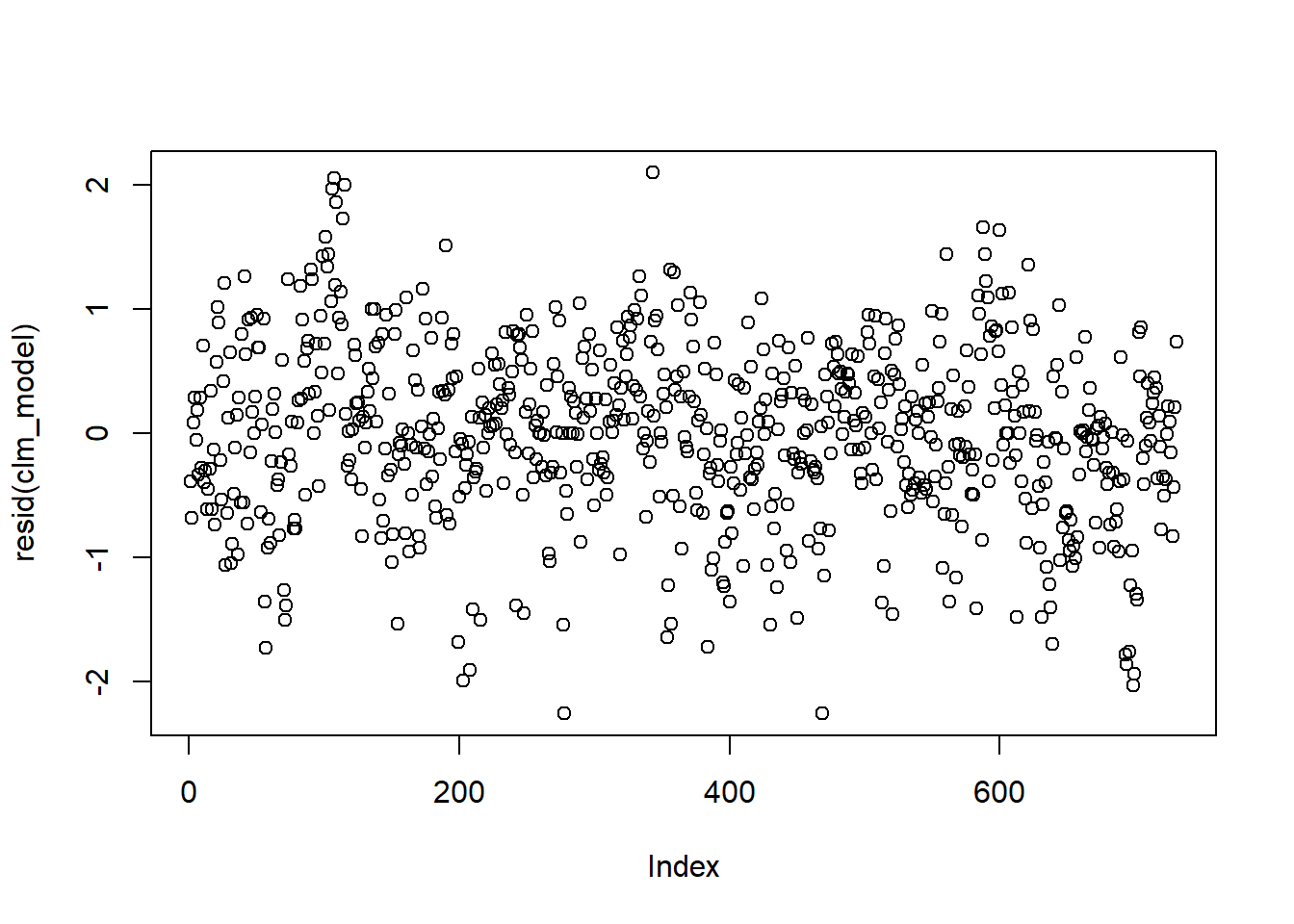

Supplementary Figure 4: Residuals of the fitted models. These residuals are normal-distributed implying good model fit.

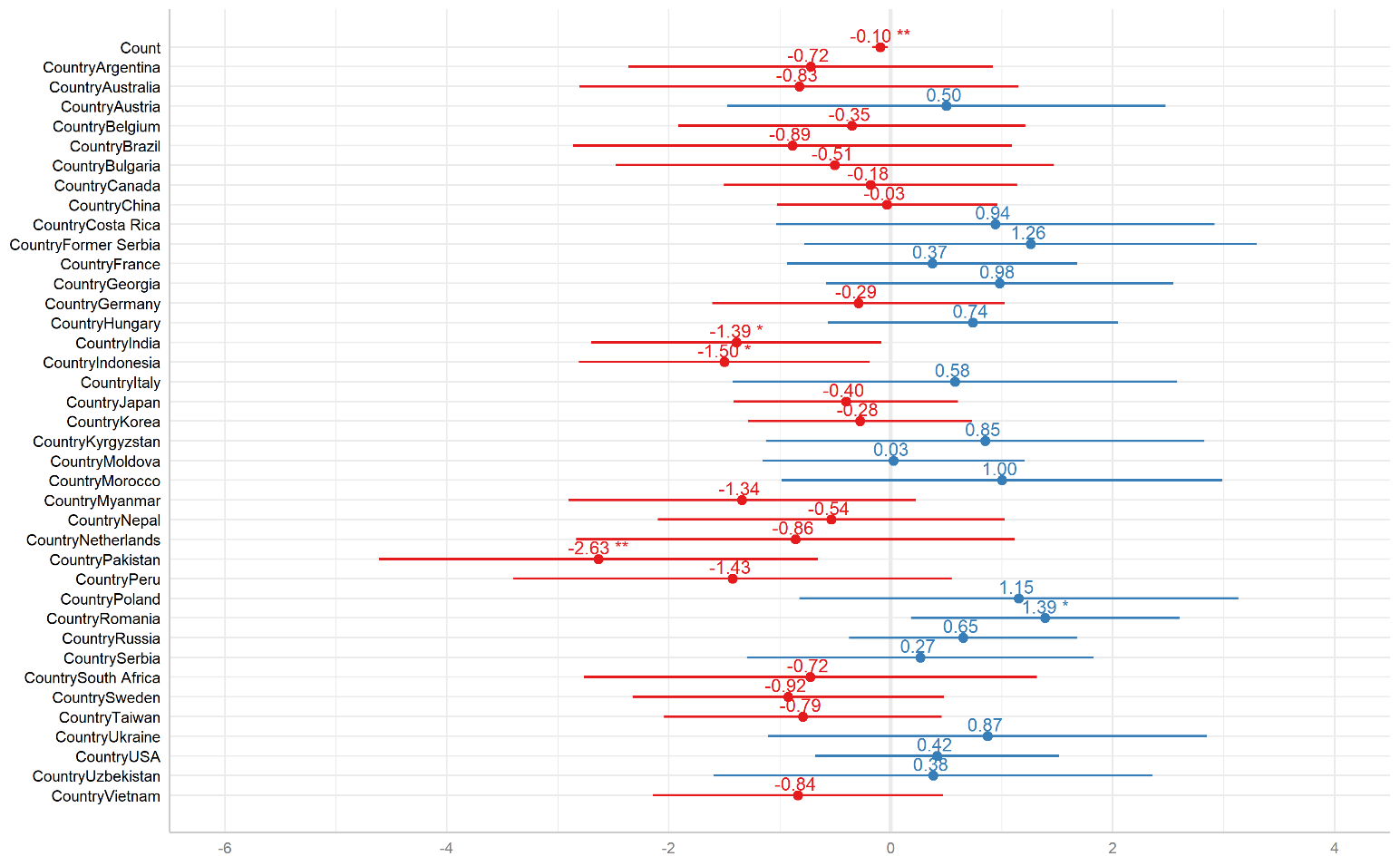

Supplementary Figure 5: Effects of all covariates country on yield prediction using landraces as baseline. Red: average effect is below zero. Blue: average effect is above zero.

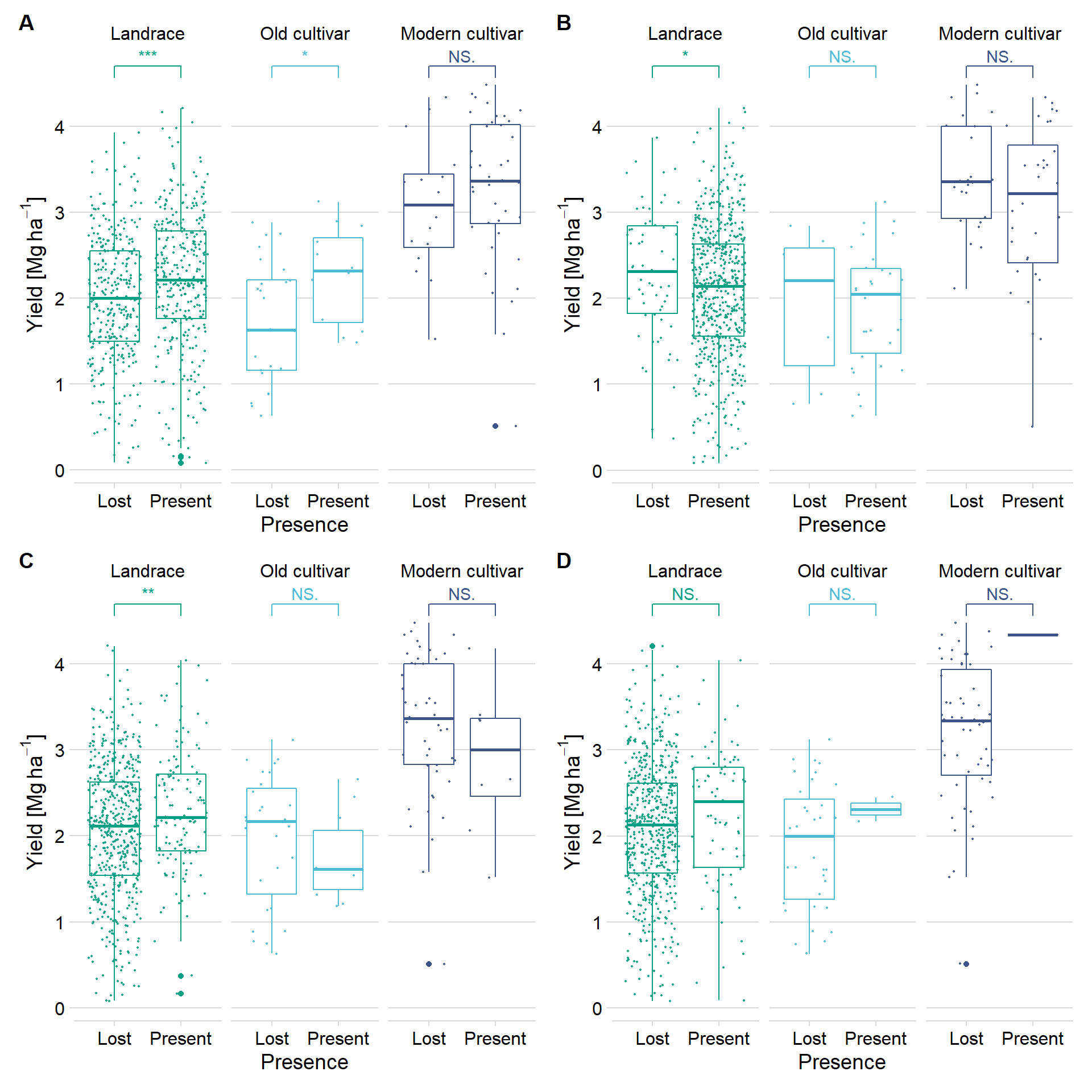

Supplementary Figure 6: Yield compared with presence/absence of four PAV-GWAS candidate genes. A) GlymaLee.01G030900 B) UWASoyPan00316 C) UWASoyPan00772 D) UWASoyPan04354. NS: Not significant, *** p < 0.001, **: p < 0.01, *: P <0.05, Wilcoxon Rank Sum Test).

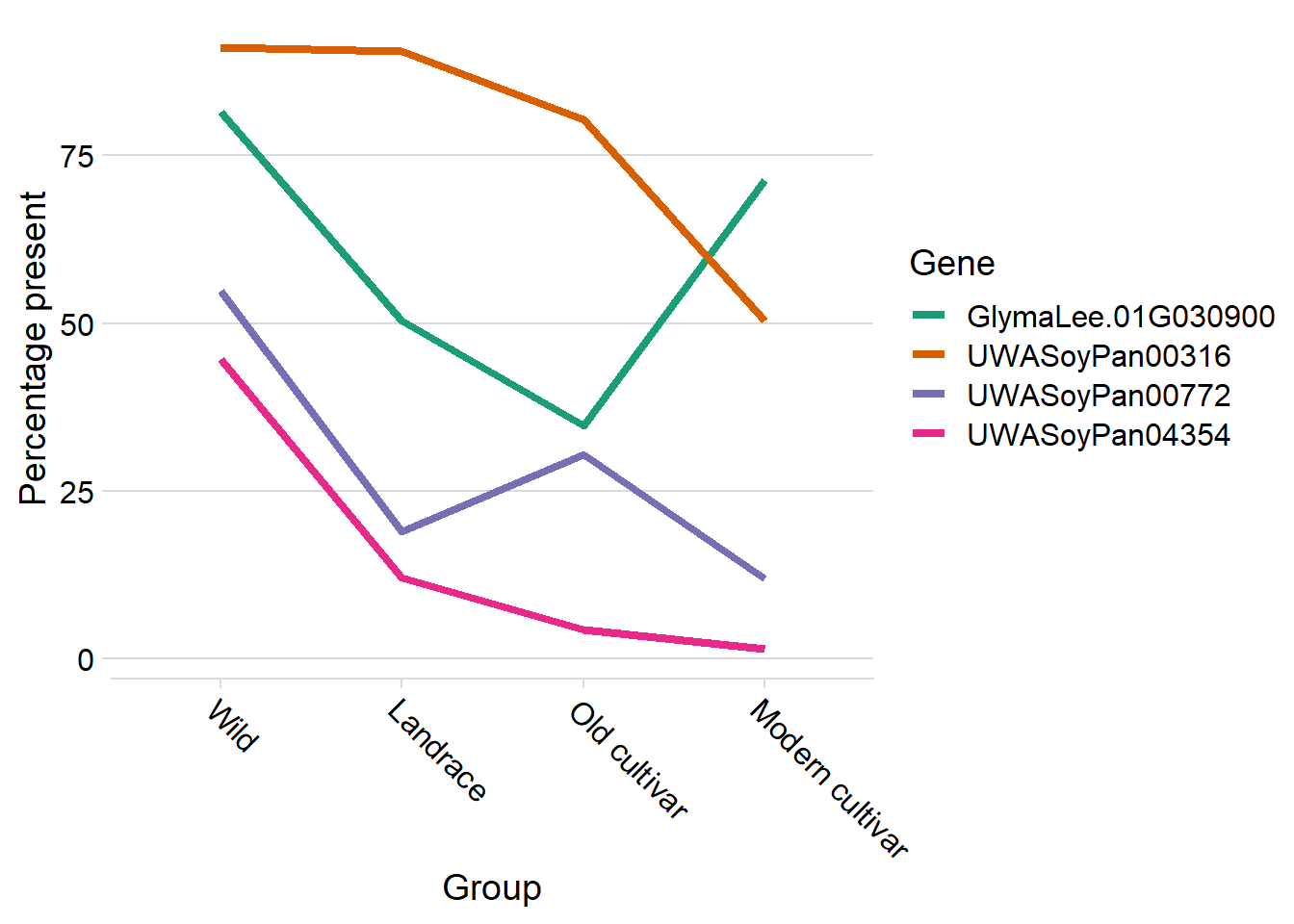

Supplementary Figure 7: Presence of four PAV-GWAS candidate genes across the four accession groups showing a generally negative trend during the history of soybean breeding, except for GlymaLee.01G030900 which increased in frequency in modern breeding line.

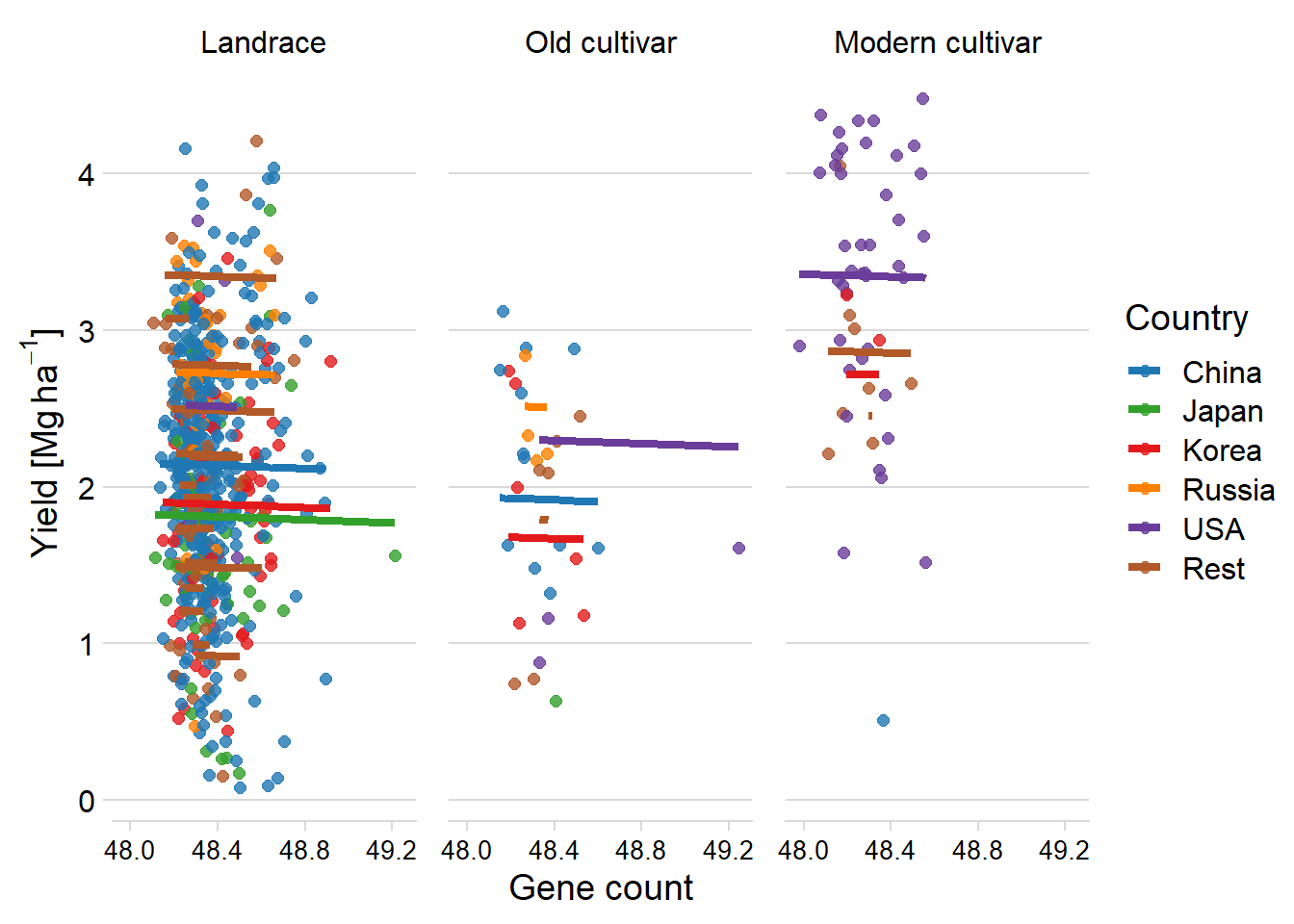

Supplementary Figure 8: Modelling of yield based on the number of all genes per line using country and accession group showing no statistically significant association between yield and the number of NLR genes. The plot is coloured by country with the five most-common countries labelled. Gene count in thousands.
